# Genetic coupling of signal and preference facilitates sexual isolation during rapid speciation

**DOI:** 10.1101/694497

**Authors:** Mingzi Xu, Kerry L. Shaw

**Affiliations:** Department of Neurobiology and Behavior, Cornell University, 215 Tower Rd, Ithaca, NY 14853

## Abstract

The divergence of sexual signals is ultimately a coevolutionary process: while signals and preferences diverge between lineages, they must remain coordinated within lineages for matings to occur. Divergence in sexual signals makes a major contribution to evolving species barriers. Therefore, the genetic architecture underlying signal-preference coevolution is essential to understanding speciation but remains largely unknown. In *Laupala* crickets where male song pulse rate and female pulse rate preference have coevolved repeatedly and rapidly, we tested two contrasting hypotheses for the genetic architecture underlying signal-preference coevolution: linkage disequilibrium between unlinked loci and genetic coupling (pleiotropy of a shared locus or tight physical linkage). Through selective introgression and quantitative trait locus (QTL) fine mapping, we estimated the location of QTL underlying interspecific variation in both female preference and male pulse rate from the same mapping populations. Remarkably, map estimates of the pulse rate and preference loci are 0.06-0.23 cM apart, the strongest evidence to date for genetic coupling between signal and preference loci. As the second pair of co-localizing signal and preference loci in the *Laupala* genome, our finding supports an intriguing pattern, pointing to a major role for genetic coupling in the quantitative evolution of a reproductive barrier and rapid speciation in *Laupala*. Due to its effect on suppressing recombination, a coupled, quantitative genetic architecture offers a powerful and parsimonious genetic mechanism for signal-preference coevolution and the establishment of positive genetic covariance on which the Fisherian runaway process of sexual selection relies.

## Introduction

From the courtship dances of birds of paradise to the songs of crickets, species commonly differ in courtship behaviors^1–3^. Because variation in sexual signals and the associated preferential responses can ultimately give rise to reproductive barriers between species, divergence of sexual signaling systems may be a potent driving force of speciation^3–5^. Hidden in the divergence of sexual signaling systems is a coevolutionary process: while signals and preferences diverge among lineages, they are functionally constrained to maintain effective communication, and thus, to coevolve within a lineage^6,7^. What genetic architecture facilitates signal-preference coevolution? Because response to selection depends on the underlying genetics, the answer to this question is indispensible to understanding the evolution of sexual signaling systems and speciation.

Two contrasting hypotheses for the genetic architecture underlying signal-preference coevolution have been proposed. The first hypothesis posits that genetic variation in sexual traits and preferences is caused by unlinked loci. Coevolution is mediated through preferential mating that results in linkage disequilibrium between trait and preference alleles over time^8,9^. In contrast, the second hypothesis proposes that genetic coupling (a shared, pleiotropic locus or tightly linked sexual trait and preference loci) underlies variation in both sexual traits and preferences^10–14^. Particularly under pleiotropy, genetic covariance is realized by mutations that affect both traits simultaneously, enhancing the efficacy of divergent selection on the communication system. Under both pleiotropy and tight physical linkage, divergent signal-preference systems are resistant to the homogenizing effects of gene flow when species hybridize because recombination between signal and preference alleles is suppressed. In contrast, under the first hypothesis, recombination can decrease the genetic covariance between unlinked signal and preference loci and even reverse speciation. The two genetic architectures thus may differ in their potency in promoting and maintaining speciation.

Theoretical models of sexual selection often assume that signal and preference loci are unlinked and that a positive genetic correlation between these traits arises through assortative mating^15–22^. Indeed, genetic mapping studies have provided support for the hypothesis of unlinked loci in chemical, acoustic and visual signaling modalities^23–25^. In contrast, genetic coupling are often considered unlikely^19^. However, recent evidence from quantitative trait locus (QTL) mapping and introgression studies supports the presence of colocalized genes underlying interspecific signal-preference variation in crickets, butterflies, fruit flies, and fish^26–30^. In addition, lab-induced mutations that alter both male signals and female preferences in fish and flies^31–33^ demonstrate that pleiotropic genes underlying signals and preferences do exist in the genomes of sexual organisms.

The Hawaiian cricket *Laupala* presents a powerful system to investigate the genetic architecture underlying signal-preference coevolution during divergence in sexual signaling systems. Rapid speciation in this genus has resulted in 38 morphologically and ecologically similar species distinguished by marked differences in acoustic behavior ^34^. Both male song and female acoustic preference have diverged repeatedly between, but remain coordinated within, species^34–37^. Like most crickets, *Laupala* males sing rhythmic songs that attract females^7,38^. Moreover, female preference for the salient feature, pulse rate, can be studied in computer playback experiments wherein females indicate preferences by phonotaxis (i.e., orienting and walking towards the preferred song). Thus, female preference for pulse rate can be easily isolated and measured. In addition, variation in acoustic behaviors both within and between species is quantitative^29,36,37^, exemplifying a common form of trait evolution in natural systems. Finally, acoustically distinct species of *Laupala* can be hybridized, allowing genetic analysis of natural variation in acoustic behavior.

In support of the genetic coupling hypothesis, a previous study of the fast singing *L. kohalensis* (pulse rate 3.72 pulse per second, pps) and the slow singing *L. paranigra* (pulse rate: 0.71 pps) demonstrated a shared QTL underlying song and preference variation on linkage group one (LG1). These co-localized loci explain approximately 9% and 15% of the species difference in pulse rate and pulse rate preference, respectively. We subsequently isolated and fine-mapped a song QTL on LG5 that explains an additional ~11% of the pulse rate difference of this species pair^39^. Marker association studies have predicted the existence of a preference QTL on LG5, yet its location on this linkage group remains unknown^40,41^.

Here, we present the remarkable discovery of a second QTL for female acoustic preference, whose map position coincides with the male pulse rate QTL on LG5. This represents the second case of shared QTL underlying signal-preference coevolution in *Laupala*. This finding illuminates the quantitative dynamics of co-evolution in sex-limited traits under sexual selection. Further, our results suggest that genetic coupling may be more common and important than we previously thought in the evolution of signal-preference sexual communication systems.

## Results

We obtained peak preference measures from 56 and 33 F_2_ females in 4C.9 and 4E.1 respectively, and 21 females in the parental near isogenic line 4C (NIL4C) and *L. kohaensis* lines. As expected, females from the control *L. kohalensis* line preferred fast pulse rate (3.78 ± 0.10 pps, mean ± SD, n = 17) and females from NIL4C preferred slow pulse rate (3.34 ± 0.05 pps, n = 4, Fig. 1).

**Fig. 1.**
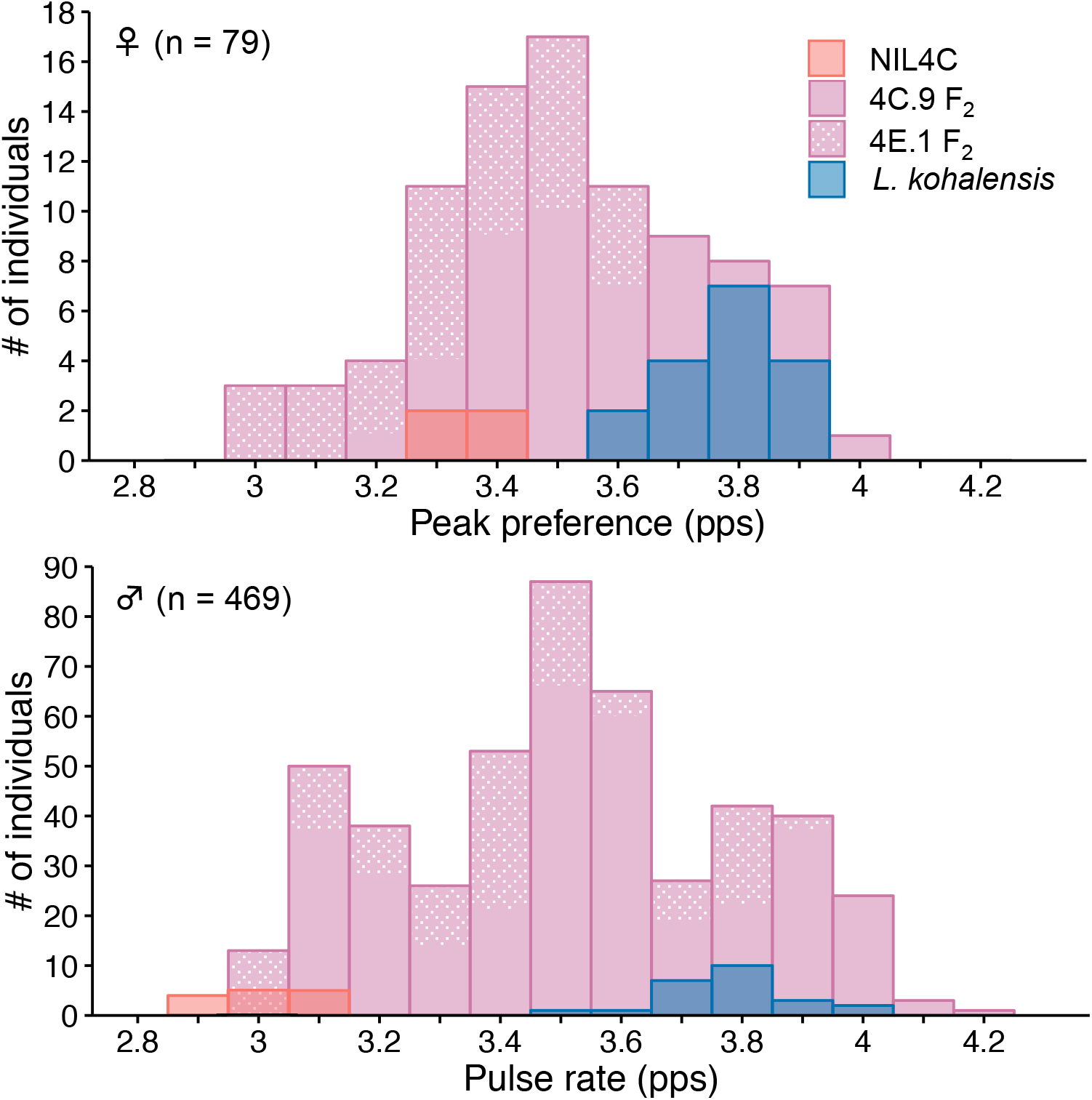
Phenotypic distributions of males and females in the parental *Laupala kohalensis*, near isogenic line 4C (NIL4C), and two F_2_ mapping families 4C.9 and 4E.1.

Using multiple imputation (IMP), we localized a preference QTL from combined 4C.9 and 4E.1 families that explained 58.8% of F_2_ variance in female preference to a peak in the LOD profile between 26.17 and 26.34 cM (Table 1, Fig. 2) on the integrated LG5 (Supplementary results, Supplementary Fig 1,). The final multiple QTL mapping (MQM) model detected a single QTL at the same location (Supplementary Fig. 2). The 1.5-LOD confidence interval spanned 2.79 cM (Table 2) and included 8 SNP markers from 7 scaffolds (Fig. 2). Three markers on scaffold S001371, S006506, and S000353 had the highest LOD score among all markers (Fig. 2).

**Table 1.** Results from multiple QTL mapping for variation in male song pulse rate and female peak preference for pulse rate using combined data from two F_2_ mapping families. Only results for the major-effect pulse rate QTL is shown in this table

| Trait | Sex | Sample size | QTL location (cM) | LOD score (LOD threshold) | 1.5-LOD CI (cM) | QTL effect size --additive (pulse / s) | % species difference explained | % F<sub>2</sub> variance explained |
| --- | --- | --- | --- | --- | --- | --- | --- | --- |
| Preference | Female | 71 | 26.17-26.34 | 7.35 (2.63) | 25.20-27.99 | 0.13 ± 0.03 | 4.32 | 58.76 |
| Pulse rate | Male | 450 | 26.40 | 207.09 (2.51) | 26.34-26.80 | 0.33 ± 0.01 | 10.96 | 85.61 |

**Fig. 2.**
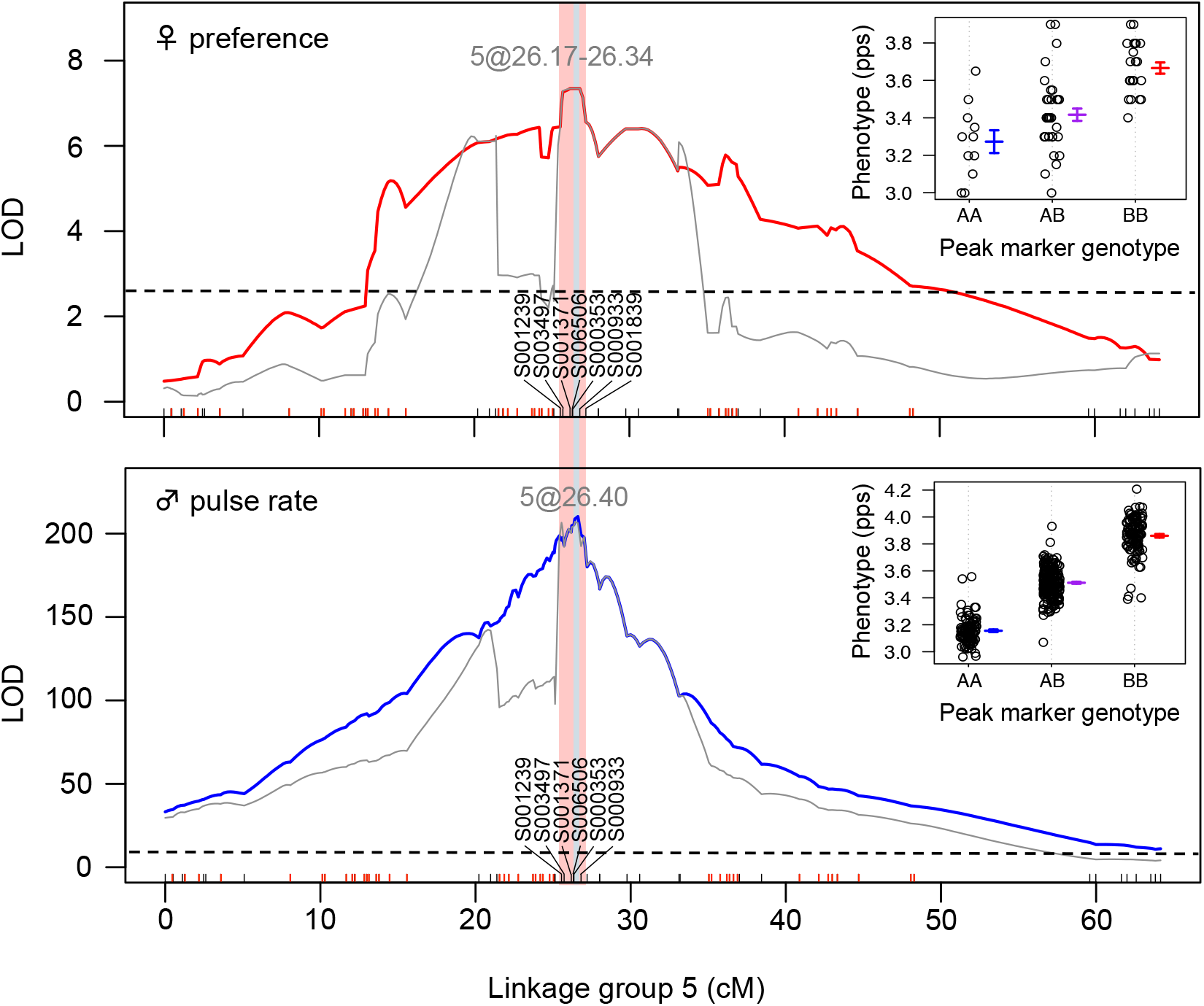
LOD profiles and phenotypic effect of alleles at the markers with the highest LOD score from the multiple imputation (IMP) models for interspecific variation in pulse rate and preference. LOD profiles from IMP models using genotypes of markers segregating at 1:2:1 ratio in both families (genotypes at markers segregating in only one family were treated as missing data, and thus simulated, in the non-segregating family) were shown in red (female preference) and blue (male pulse rate). LOD profiles from IMP models using all genotypes at all markers, including those not segregating in one of the two families were shown in grey. Markers (tick marks on the x-axis) marked in red indicate those segregating in one of the two families and markers in black are those segregating in both families (see Supplementary information for details). The red and blue shaded areas indicate 1.5-LOD confidence intervals for preference and pulse rate QTL respectively. The scatter plots show individual phenotypes for three genotypes at the marker with the highest LOD score for preference and pulse rate respectively.

At the markers with the highest LOD score, females with the *L. paranigra*-origin (AA), the heterozygous, and the *L. kohalensis*-origin (BB) genotype showed preference for slow (3.27 ± 0.05 pps, mean ± SE, Fig. 2), intermediate (3.42 ± 0.03 pps), and fast pulse rates (3.66 ± 0.04 pps) respectively. The phenotypic effect of a single allele at the preference QTL was largely additive (0.13 ± 0.03 pps), explaining 4.3% of the phenotypic difference between the pure species parents (Table 1). Combining 4C.9 and 4E.1, the phenotypic distribution of the female peak preference in the F_2_ generation was consistent with a 1:2:1 segregation ratio (bin1 = 21, bin2 = 45, bin3 = 23, X^2^ = 0.10, df = 2, p = 0.95, Fig. 1).

We used pulse rate measures from 339 and 130 males in 4C.9 and 4E.1 respectively (published previously in ^39^). A major-effect QTL explaining 85.6% F_2_ variance in pulse rate was localized at 26.40 cM in both IMP and MQM (Fig. 2, Supplementary Fig. 2, Table 1), consistent with previous results ^39^. The 1.5-LOD confidence interval of this QTL spanned 0.46 cM (Table 1, Fig. 2). MQM identified two additional small-effect QTL that explained 1.17% and 0.48% of F_2_ variance at 5.6 cM and 59.8 cM respectively (Supplementary Fig. 2, Supplementary Table 1).

Similar to the preference QTL, males with homozygous *L. paranigra* (AA), heterozygous (AB) and homozygous *L. kohalensis* genotype (BB) at the marker with the highest LOD score had slow (3.16 ± 0.01 pps), intermediate (3.51 ± 0.01 pps) and fast pulse rates (3.87 ± 0.01 pps, Fig. 2) respectively. The phenotypic effect of an allele at the pulse rate QTL was almost entirely additive (0.33 ± 0.01 pps), explaining 11.0% of species difference (Table 1). The F_2_ phenotypic distribution of male pulse rate was consistent with a 1:2:1 segregation ratio as shown previously ^39^ and did not significantly differ from that of female preference in F_2_ (X^2^ = 2.15, df = 2, p = 0.34).

## Discussion

Sexual signals and preferences commonly differ between species, reflecting the powerful role of signal and preference divergence in the speciation process^5,42^. While sexual traits routinely diverge, sexual selection likely constrains sexual signals and preferences to remain coordinated during this process. The genetic architecture underlying signal-preference coevolution is central to understanding the complex selective landscape underlying speciation. Signal and preference traits are often modeled with independent hereditary bases where assortative mating alone results in linkage disequilibrium among unlinked loci. Yet recent findings consistent with a coupled basis to natural variation in signals and preferences in multiple taxa^26–30^ suggest a key to understanding signal-preference coevolution. *Laupala* crickets exemplify these patterns, exhibiting distinctive male songs and female acoustic preferences that have diverged between closely related species in an extremely rapid speciation context^34^. *Laupala* are further distinct from other systems in that signal and preference are sex-limited traits involved in speciation by sexual selection. Moreover, unlike traits with simple genetic switches, both song and preference are complex, and vary quantitatively, representing a common mode of trait evolution. *Laupala* thus offers the potential for novel insights.

A significant limitation for many quantitative preference phenotypes is the ability to estimate preference in a segregating population. The relatively simple acoustic behaviors in *Laupala* allowed us to fine map a new female preference QTL on LG5 between the fast singing (and preferring) *L. kohalensis* and slow singing *L. paranigra*. We localized the preference QTL on a 0.17 cM-wide peak with a 1.5-LOD confidence interval of only 2.8 cM. Our study is one of only three to have mapped the location of preference/mate choice loci with sufficiently high resolution to rigorously test alternative hypotheses of genetic architecture^27,28^. Moreover, our study is unique among these in that acoustic preference in *Laupala* is a sex-limited, quantitative trait expressed in the context of sexual selection by female mate choice, the leading causal explanation for the evolution of elaborate sexual communication^34^.

Remarkably, we found that the estimated map position of the female preference QTL on LG5 is nearly identical (only 0.06-0.23 cM apart) to a male pulse rate QTL (Fig. 2). Furthermore, the 1.5 LOD confidence intervals of the preference and song QTL largely overlap (Fig. 2, Supplementary Fig. 2) with the top three LOD scores for preference and song attributed to the same markers. Coincidentally, both preference and song QTL contribute relatively similar magnitudes of effect in a largely additive way to the differences in acoustic behaviors between the two species (Table 2). Equally importantly, the phenotypic effects of pulse rate and preference QTL are in the same direction, required for establishing a positive genetic covariance between pulse rate and preference. Features of genetic architecture such as these can greatly facilitate the coevolution of a signal-preference system, whereby both traits vary in quantitatively small steps in the same direction, enabling coordinated changes despite the divergent phenotypic evolution that must occur during the speciation process.

The QTL identified in the present study comprise the second pair of colocalizing song and preference QTL identified in *Laupala*. We have previously mapped QTL that make a similarly coupled contribution to pulse rate and preference differences between *L. paranigra* and *L. kohalensis* on LG1^29^. At least eight and four QTL underlie the species difference in pulse rate and preference respectively between these species^29,40,41,43^. The fact that two of the four preference QTL independently coincide with different pulse rate QTL (the location of the remaining two as yet unknown) suggests a compelling pattern underlying the co-evolution of these traits. Moreover, the allelic effect of the two preference QTL and of their colocalizing pulse rate QTL together account for roughly 20% of species difference (Table 1 herein; Table 1 in^29^). Such a substantial proportion attests to a significant role that genetic coupling plays in sexual signal divergence and speciation in *Laupala*.

The overlap between the confidence intervals of the pulse rate and preference QTL and the extremely close estimates of the peak locations are consistent with a pleiotropic basis to variation in pulse rate and preference. Pleiotropy provides a genetic mechanism whereby positive genetic covariance between signal and preference genes is an immediate consequence of mutations at the locus (or loci for quantitative traits). Alternatively, our results may reflect the genetic architecture of a tightly linked signal-preference “super gene”, an equally exciting explanation that has been repeatedly shown to facilitate adaptation in complex trait suites^44–46^. Like pleiotropy, tight physical linkage can facilitate coevolution by effectively suppressing recombination between signal and preference alleles. Distinguishing pleiotropy and tight linkage requires identifying the causal genes and is a logical next step in this system. Recent technological advances place such a goal within reach in non-model systems^47^.

Mechanistically, recent findings suggest that shared genes for singing and temporal auditory pattern recognition are plausible. Insect singing by wing movements is controlled by central pattern generators (CPG) in thoracic and abdominal ganglia^48,49^. In the field cricket, an auditory feature detector circuit that selectively responds to pulse rate of conspecific song has been identified in the female brain^50^. In this circuit, pulse rate selectivity is achieved via postinhibitory rebound that offsets direct and delayed line inputs to a coincidence detector neuron by the exact duration of the conspecific pulse period. We suggest that a shared molecular mechanism, for example, the type or number of ion channels or neural projections, may regulate both the oscillation period of the song CPG and the offset duration of the feature detector circuit. Fine mapping and gene annotation^39^ identified a promising candidate gene for song variation, the putative *Laupala* cyclic nucleotide-gated ion channel-like gene (*Cngl*) on scaffold S001371. Here we show that the highest LOD score for preference also associates with this scaffold. Furthermore, a non-synonymous SNP differentiating the two parental species was found in the conserved cyclic nucleotide binding domain ^39^. Although unknown in *Laupala*, *Drosophila Cngl* is expressed in brain, thoracic ganglia and muscles ^51^, consistent with the expectation for a causal gene for song and preference variation. Finally, a related group of genes in the same gene superfamily are implicated in both rhythmic muscle contraction^52–55^ and temporal coincidence detection and relay in auditory systems^56–58^. Such evidence renders *Cngl* a candidate pleiotropic gene for further functional validation.

Colocalization of sexual traits and mate choice has been shown in two other high-resolution mapping studies. In the threespine stickleback, QTL for mate choice and body shape were 14.3 cM apart^26^. In the *Heliconius* butterflies, a QTL contributing to courtship time is only 1.2 cM from *optix*, a gene regulating the forewing red band^28^, demonstrating genetic coupling underlying variation in wing coloration and courtship time. In both these systems, sexual signals are likely magic traits^59^ that function in both ecological (foraging or predator avoidance) and mate choice contexts. In contrast, the sexual traits we have studied in *Laupala* are sex-limited and function exclusively in a reproductive context, representing the widespread process of sexual selection by female choice thought to underlie the evolution of many elaborate and extravagant sexual signaling systems. Intriguingly, the Fisherian runaway process of sexual selection is a primary explanation for the evolution of exaggerated sexual traits that relies on positive genetic covariance between sexual trait and preference^8,19^. Whether such positive genetic covariances exist is debated^60,61^; our finding offers a parsimonious and effective genetic mechanism for the establishment and maintenance of positive genetic covariance for the trait pair^19,22,62^.

Taken together with the studies above, genetic coupling may transcend communication modality, evolutionary mechanism and taxonomic group (invertebrates or vertebrates) and prove to be of general importance to the divergence of sexual communication systems and speciation. In light of these emerging empirical findings, theoretical and modeling work has recently begun to explore the effect of pleiotropy and tight physical linkage on the process and consequence of lineage divergence^63,64^. Further empirical and theoretical efforts should be fruitful in deepening our understanding of how divergence in communication systems can spur speciation.

## Methods

### Breeding design

Details of the breeding design have been described in^39^. Briefly, through selective introgression, we isolated and introgressed the largest-effect QTL for male song pulse rate (QTL4) from *L. paranigra* into the *L. kohalensis* genome in two independent near isogenic lines (NIL4C and NIL4E)^40,65^. NIL males were then backcrossed with *L. kohalensis* females to generate two F_2_ mapping families for two NILs (denoted as family 4C.9 and 4E.1). All crickets were reared individually in 120 ml specimen cups with a piece of moist tissue and fed *ad lib* Organix organic chicken and brown rice dry cat food (Castor & Pollux Natural Petworks, Clackamas, USA) twice per week at 20.0°C and light cycle at 12h:12h light:dark.

### Male song simulation

We simulated digital songs for female preference trials in LabVIEW v.8.2^66^. The simulated songs consisted of pulsed sinusoidal tones with characteristics of natural *Laupala* songs (40 ms pulse duration, 5 kHz carrier frequency, also see Supplementary methods). In a preference trial, songs played simultaneously from two speakers were calibrated to 90 dB at the cricket release point, 75 cm from the speaker (i.e., at the center of the phonotaxis tube, see below).

### Phenotyping

Methods for phenotyping male song pulse rate were reported in^39^. Here, we measured the peak preference for pulse rate from females using repeated, two-choice phonotactic trials. The trials were conducted in custom-made phonotaxis tubes (Fig. 3a, also see Supplementary methods) in a RS-243 ETS-Lindgren's sound isolation booth (ETS-Lindgren, Wood Dale, USA) at 20 °C.

**Fig. 3.**
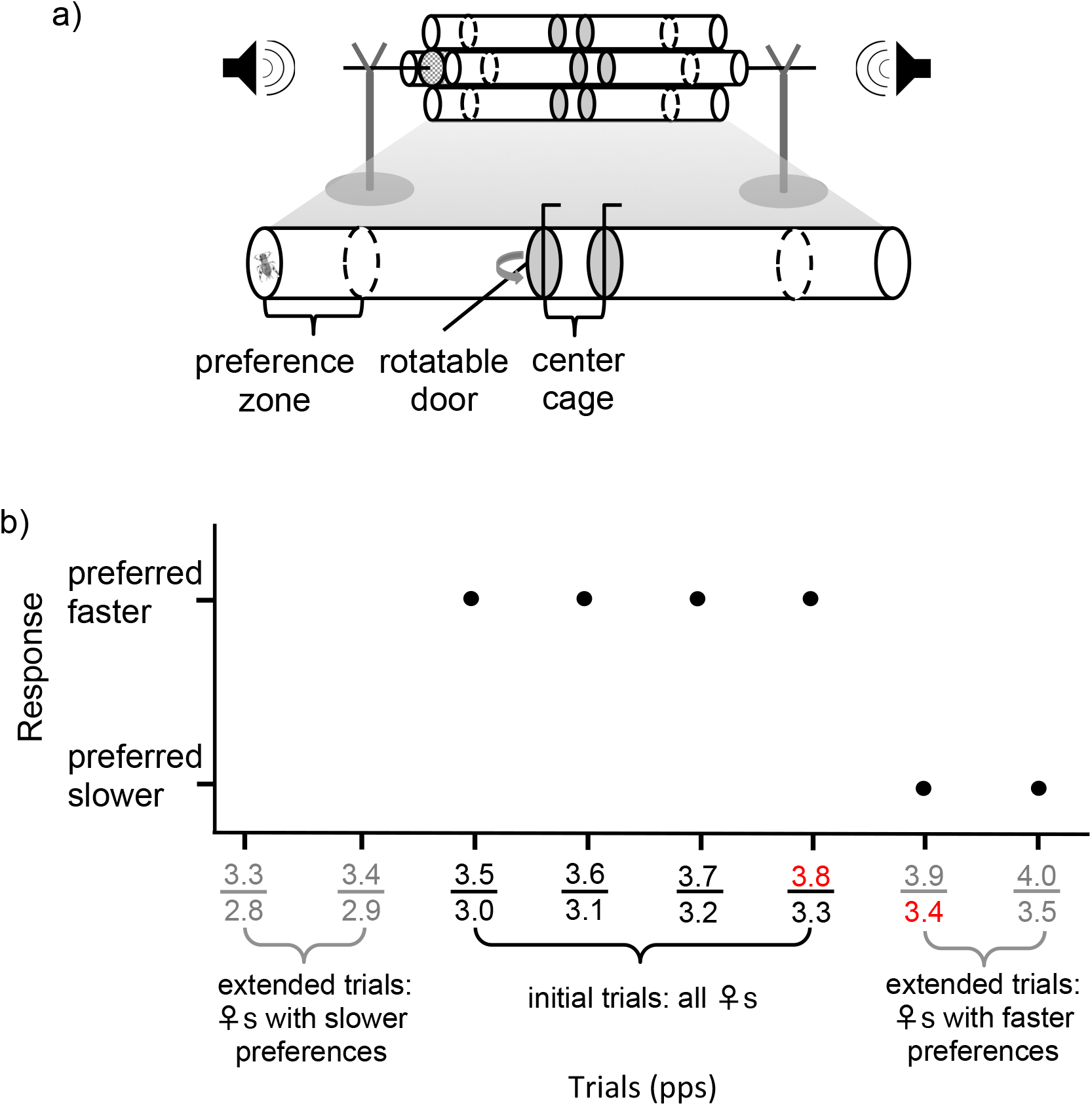
a) Preference trial apparatus and b) experimental design of the preference trials, response data from a hypothetical female that goes through 4 initial trials and 2 extended trials at the faster end, and inference of peak preference. The pulse rates of the two song played at a given trial were given on the x-axis with the faster pulse rate above and the slower pulse rate below. In the example here, the female shows a switch from preferring the faster pulse rate to preferring the slower pulse rate between trial 3.3 v.s. 3.8 pulse/s (pps) and trial 3.4 v.s. 3.9 pps. The peak preference is thus calculated as (3.8+3.4)/2=3.6 pps.

Each phonotaxis trial consisted of a 5-min pretrial period and a 10-min testing period. Two simulated songs differing by 0.5 pps but were otherwise identical were broadcasted simultaneously during both the pretrial and testing periods from speakers placed 180° apart (Fig. 3a). Songs were randomized by speaker for each trial. During the pretrial period, the focal female was confined to the central cage. To commence a trial, the doors at both ends of the central cage were opened to connect the cage with the phonotaxis tube. If the focal female entered the preference zone defined as the last 10 cm at each end of the tubes, we scored a preference for the song pulse rate from that speaker.

Each female was tested in a series of trials to estimate peak preference (Fig. 3b). All females were initially tested in 4 trials in random order where the pulse rates were 3.2 v.s. 3.7 pps, 3.3 v.s. 3.8 pps, 3.4 v.s. 3.9 pps, and 3.5 v.s. 4.0 pps for 4C.9, and 3.0 v.s. 3.5 pps, 3.1 v.s. 3.6 pps, 3.2 v.s. 3.7 pps, and 3.3 v.s. 3.8 pps for 4E.1. The pulse rate range of the initial four trials was determined by the F_1_ male pulse rate distribution in each family. If female response switched from faster pulse rates at the lower end to slower pulse rates at the higher end of the trial range, her peak preference was estimated on the basis of these four trials as the midpoint of the switch from faster to slower pulse rates (Fig. 3b). If the female showed consistent response to either faster or slower pulse rates in the initial four trials, she was further tested in extended trials at either the lower (4C.9: 3.0 v.s. 3.5 pps, 3.1 v.s. 3.6 pps; 4E.1: 2.8 v.s. 3.3 pps and 2.9 v.s. 3.4 pps) or higher (4C.9: 3.6 v.s. 4.1 pps and 3.7 v.s. 4.2 pps; 4E.1: 3.4 v.s. 3.9 pps and 3.5 v.s. 4.0 pps) end of the range, depending on the direction of her response in the initial trials (Fig. 3b). We repeated each trial up to three times for females who failed to respond in a given trial. On any given day, females were tested in no more than two trials, with at least 2h between the trials. In cases where a female consistently showed preference for faster or slower pulse rates in all 6 trials, we estimated the peak preference at the most conservative value (i.e., the midpoint in the next extreme trial, assuming the female would show a switch in her preference).

### Genotyping and linkage mapping

Genotyping has been reported in^39^. Briefly, we sequenced F_2_ individuals using Genotyping-by-Sequencing^67^. Genotypes of single nucleotide polymorphism (SNP) markers were called in each family using the *L. kohalensis* genome reference as the *L. kohalensis* parent (whose genotype was denoted as “B”); the alternative allele (denoted as “A”) was assigned to the NIL parent. We excluded SNPs that deviated significantly from a 1:2:1 segregation ratio (Benjamini-Hochberg adjusted p < 0.05) and had a mean depth of coverage < 20 in each family. Using the resulting SNPs, linkage maps of LG5 for family 4C.9 and 4E.1 have been constructed previously^39^ and integrated herein in Joinmap 4^68^. We kept one SNP marker per 2 kb on the same scaffold. When markers on the same scaffold showed order conflict between the two families, we removed the marker with fewer unique pairwise recombinations until the marker order conflict is resolved.

### QTL mapping

QTL mapping for pulse rate in each family was performed previously^39^. Here, we combined data from both families to increase power in QTL mapping of female peak preference on the integrated map. We also remapped the pulse rate QTL using combined data on the integrated map. Individuals with < 25% missing genotypes were used for QTL mapping. We first tested for a family effect on phenotype using Welch's t test. We then performed multiple imputation (IMP) with family as an additive covariate. As a minor-effect QTL for song pulse rate has been detected previously, we also performed multiple QTL mapping (MQM) for both pulse rate and preference in case the existence of other minor-effect QTL affects location and effect size estimation of the focal QTL. Missing genotypes were simulated by 10000 multiple imputations. LOD thresholds were calculated from 1000 permutations at an α level of 0.05. We estimated effect sizes of significant QTL by both the final MQM models and by phenotypes at the marker with the highest LOD score. All QTL mapping analyses were conducted in R/qtl v.1.39-5 (Broman et al. 2003).

Finally, we tested whether the phenotypic distribution of female preference in the combined F_2_ families deviated from a 1:2:1 Mendelian segregation ratio with a two-sided chi-square test. To do so, we binned phenotypic data by dividing the range of the phenotypic values evenly in three bins.

## Data availability

DNA sequences are available at NCBI (accession PRJNA509479). Phenotype and genotype data as well as Bioinformatic, QTL mapping and LabVIEW scripts can be downloaded at https://github.com/MingziXu/QTL4-colocalization-scripts.

## Acknowledgements

We thank Ben Weaver, Alex Thomas, Eric Cole, Laura Hernandez, Isabelle Phillipe, Nathan Barr and McKenzie Laws for assistance in cricket rearing, phenotyping, and DNA extraction, Cornell Genomic Diversity Facility for advice on molecular procedures, and Aure Bombarely, Thomas Blankers, Linlin Zhang and Cornell BioHPC lab for assistance in bioinformatics. This project is funded by National Science Foundation grant #1257682 to KLS.

## Notes

https://github.com/MingziXu/QTL4-colocalization-scripts

https://www.ncbi.nlm.nih.gov/search/all/?term=PRJNA509479

