## Supplemental Information for "Genetic coupling of signal and preference facilitates sexual isolation during rapid speciation"

**Supplemental Methods**

Male song simulation

In the simulated song, each pulse had a duration of 40 ms, a carrier frequency of 5 kHz, and an amplitude envelope of 10 ms rise and 30 ms fall period. These song parameters are characteristic of the natural songs and do not differ significantly between the two species.

Sound broadcasting system

In female preference trials, simulated digital songs from LabVIEW were passed to a RM1 mobile processor to generate a two-channel analog output (Tucker-Davis Technologies, Alachua, USA), one for each song. The analogs were in turn, passed through a Krohn-Hite frequency filter model 3322 (Krohn-Hite Corporation, Brockton, USA) to filter out frequencies higher than 6 kHz and a Crown D-75A amplifier (Crown Audio, Elkhart, USA). The songs were then broadcasted by two RadioShack speakers (model 40-1218, RadioShack, Fort Worth, USA) in a RS-243 ETS-Lindgren's audiometric exam booth (ETS-Lindgren, Wood Dale, USA). Before trials each day, we calibrate the speakers to 90 dB 75 cm from the speaker (i.e., at the center of the phonotaxis tube) using a Brüel and Kjær sound level meter (model 4155, Brüel and Kjær Sound and Vibration Measurement, Nærum, Denmark).

Phonotaxis apparatus design

The phonotaxis tubes were 90 cm in length and 8 cm in diameter and were made of cardboard frames and black window screen walls. A removable, cylindrical cage 15 cm long and 8 cm in diameter was mounted in the center of the tube. The cage had two screen doors to connect with the phonotaxis tubes. Four phonotaxis tubes were arranged on a central axis with equal space between them such that four females can be tested simultaneously. Two speakers were placed at the same level of the center axis (and hence, at equal distance to each of the four tubes) and are 75 cm from the center of the cages.

Genotyping

In the first step of genotyping, within each family, we called SNPs using FreeBayes v.0.9.12-2-ga830efd (61) and filtered SNPs using VCFtools 0.1.15 (62) and vcffilter in vcflib v.1.0.0 (63). We retained bi-allelic SNP markers that fulfill the following criteria: 1) < 20% missing data per family; 2) minor allele frequency ≥ 2.5%; 3) genotype depth ≥ 5; 4) Phred scaled variant quality ≥ 30; and 5) strand balance probability for reference and alternative alleles > 0.0001.

The integrated linkage map includes SNPs common to both families as well as unique to each family that were non-segregating in the other family. Initially, the latter were not called as SNPs in the non-segregating families. For purposes of QTL mapping (see below), we either designated these invariant markers as “missing data” in the relevant family and simulated SNP’s based on flanking marker status (analysis A), or force-called SNP’s (analysis B) in 4C.9 at sites unique to 4E.1, and vice versa, using "--variant-input" and "--only-use-input-file" functions in FreeBayes with the minimum alternative variant count and fraction set to 0. The "--variant-input" function, when supplied with the vcf file of the unique SNPs in one family, forces FreeBayes to treat these SNPs as putative variants even if these is not enough support to pass input filters in the other family. The "--only-use-input-file" function asks FreeBayes to output variant calls and genotypes only at sites in the input vcf file. We compared the QTL results of these two strategies (see below).

Integration of LG5 maps

We integrated the linkage maps of LG5 of 4C.9 and 4E.1 using the regression algorithm with Kosambi mapping function. Specifically, we used linkages with maximum recombination frequency of 0.4 and a log-of-odd (LOD) score higher than 1, as well as a goodness-of-fit jump threshold for removal of markers at 4.

QTL mapping and effect size estimation

For both multiple imputations and multiple QTL mapping, genotype probability was calculated at a step size of 0.2 cM and genotyping error rate of 0.1% under Kosambi map function. Multiple QTL mapping was conducted using forward selection with backward elimination. Specifically, we began with a single-QTL model at the location of the significant QTL from the multiple imputation model. We then scanned for additional QTL on the integrated LG5. Significant QTL with the highest LOD score was added to the subsequent model in each round. Any additional QTL and interactions with previous QTL were accepted only when: (1) the LOD score of the main effect or the interaction term from ANOVA when each term is dropped from the model exceeded the threshold for additive or interaction term, and (2) the increase in LOD score of the overall new model compared to the previous model was greater than the penalty for the additional degrees of freedom. The penalty value controls the rate of including extraneous terms in the model at a target rate of 5%. For additive models, we used the main-effect penalty and for models with interactions, the heavy interaction penalty that controls false positive rate for models of any size was used. QTL locations were refined after each step with an iterative maximum likelihood algorithm. We repeated the process until no significant additional QTL or interaction terms were found. LOD thresholds for main and interaction terms in MQM were calculated from 1000 permutations using Haley-Knott regression. We used Haley-Knott regression instead of multiple imputations due to computational constraints.

The proportion of the phenotypic differences between the two parental species explained by a QTL was calculated as the additive effect (i.e., effect of substituting one allele) divided by the total difference in mean phenotypic values of the two parental species. When there is no dominance effect, this estimate has a maximum value of 50% of the species difference because it is a haploid effect. The proportion of $F_2$ variance explained by a QTL was calculated from ANOVA tests dropping one QTL at a time from the model. When multiple linked QTL were identified, the proportion of $F_2$ variance explained by the major QTL was estimated from a model including the major QTL only, and the proportion of $F_2$ variance explained by the minor QTL were estimated by an ANOVA test dropping the minor QTL from the final multiple QTL model.

**Results and Discussion**

Linkage mapping

The integrated SNP maps from the two $F_2$ families (4C.9 and 4E.1) in the two NIL replicates had a total length of 64.16 centiMorgan (cM) and included 75 SNP markers on 67 unique scaffolds, among which 33 markers were segregating in both families, 41 were segregating in 4C.9 only and 1 was segregating in 4E.1 only (Figure S1). The median marker interval on the linkage map is 0.41 ± 0.77 cM (median ± interquantile range).

Preference phenotype sample size

We phenotyped 393 and 112 $F_2$ females in 4C.9 and 4E.1 respectively, among which 129 and 57 females responded in at least one trial. Eleven and six females from 4C.9 and 4E.1 showed inconsistent preferences and were excluded from subsequent analyses. We obtained peak preference measures from 56 and 33 $F_2$ females in 4C.9 and 4E.1. Among these females, 14 and 10 in 4C.9 and 4E.1 did not show a switch in extended trials and peak preference was estimated by inferring a switch in the next extreme trial.

Family effect

The two $F_2$ families showed approximately equal variance (4C.9: 0.04, 4E.1: 0.03, Bartlett test, $K^2 = 0.25$, df = 1, $p = 0.62$) but differed significantly in mean peak preference (4C.9: 3.61 ± 0.20 pps, 4E.1: 3.34 ± 0.18 pps, mean ± SD, $t = 6.48$, df = 71.3, $p < 0.0001$). Therefore, family was used as a covariate in all mapping models for preference below. Likewise, family was used as a covariate in all mapping models for pulse rate as pulse rate differed significantly between the two families, and in the same direction as females (4C.9: 3.54 ± 0.28 pps, 4E.1: 3.45 ± 0.24 pps, $t = 3.88$, df = 238.2, $p = 0.0001$).

QTL mapping

We conducted QTL mapping twice, first (hereafter, analysis A) using genotypes at all marker loci wherein genotypes at marker loci segregating in only one family (Figure S1, markers in white) were treated as missing data in the non-segregating family and simulated in R/qtl; and second (hereafter, analysis B) using genotypes at all marker loci wherein genotypes at marker loci that were segregating in only one family were force-called in the non-segregating family. More detail follows, but for female preference, estimates of peak location and confidence interval did not differ between analyses A and B. For male pulse rate, the two analyses gave the same confidence interval estimation, but the peak location differed by 0.2 cM (26.60 cM in analysis A and 26.40 cM in analysis B), thus having little impact to our final conclusions.

Over the section of the integrated map where marker loci were segregating in both families (Figure 2, Figure S1, markers with checked pattern, Figure S2, black tick marks), the LOD profiles from the two analyses were exactly the same for both pulse rate and preference. This is expected because the genotypes at these marker loci in the two analyses were exactly the same. However, over the sections of the integrated map where marker loci were segregating in only one of the two families (Figure 2, Figure S2, red tick marks), the LOD profile from analysis B was below the LOD profile from analysis

A. This pattern was consistent in both IMP and MQM, and for both male pulse rate and female preference.

The lower LOD profile from analysis B at marker loci segregating in only one family was expected. Because these marker loci were homozygous in the non-segregating family (mostly in 4E.1, see Figure S1 and Supplemental Results and Discussion, Linkage mapping section), there can be no association with the observed phenotypic segregation in that family. Therefore, overall, we expect lower LOD scores for these markers than if they had been segregating in both families. In analysis A, genotypes at the marker loci in the non-segregating family were treated as missing data; R/qtl, simulates genotypes based on evidence of segregating genotypes in the flanking regions. LOD scores calculated using these simulated genotypes were consequently higher than those calculated using the actual homozygous genotypes. Therefore, the depression in LOD profile in analysis B is not mapping error or artifact, but rather, reflects stronger predictive power enabling us to rule out larger sections of the linkage map as the likely locations for the QTL peak.

A consequence of the lower LOD profile in analysis B is a relative "dip" that creates a false peak for both pulse rate and preference to the immediate left of the depressed LOD, between 21.54 and 25.20 cM in the LOD profile (Figure 2, Figure S2). This false peak was created by the adjacent depression in the LOD profile at marker loci that were segregating only in 4C.9 but were homozygous in 4E.1. Indeed, this false peak was not recognized as a significant QTL in any model for either male pulse rate or female preference.

Forty-one of the 42 markers that were segregating in only one family were homozygous in 4E.1. These non-segregating regions in 4E.1 likely resulted from multiple double recombination events occurring during previous generations of selective introgression. Consistent with this hypothesis, individuals from RIL4E line (siblings of the RIL male that generated 4E.1) all had homozygous *L. kohalensis* genotypes at marker loci in regions of lower LOD profile in analysis B.

Due to the advantage in predictive power, we report results from analysis B in the main text but LOD profiles of both analyses are shown in Figure 2 and Figure S2.

Table S1. Sample sizes, location estimations, LOD scores, confidence intervals and phenotypic effect estimations of three QTL for male song pulse rate variation from the final multiple QTL mapping using combined data from two $F_2$ mapping populations

| Trait | Sex | Sample size | Linkage group | QTL location (cM) | LOD score | 1.5-LOD CI (cM) | QTL effect size --additive (pulse / s) | % species difference explained | % $F_2$ variance explained |
|---|---|---|---|---|---|---|---|---|---|
| Pulse rate | Male | 450 | 5 | 5.60 | 10.71 | 0.00-11.60 | 0.05 ± 0.007 | 1.66 | 1.17 |
| Pulse rate | Male | 450 | 5 | 26.40 | 178.00 | 26.34-26.60 | 0.33 ± 0.01 | 10.96 | 85.61 |
| Pulse rate | Male | 450 | 5 | 59.80 | 4.55 | 47.80-62.20 | 0.01 ± 0.006 | 0.47 | 0.48 |

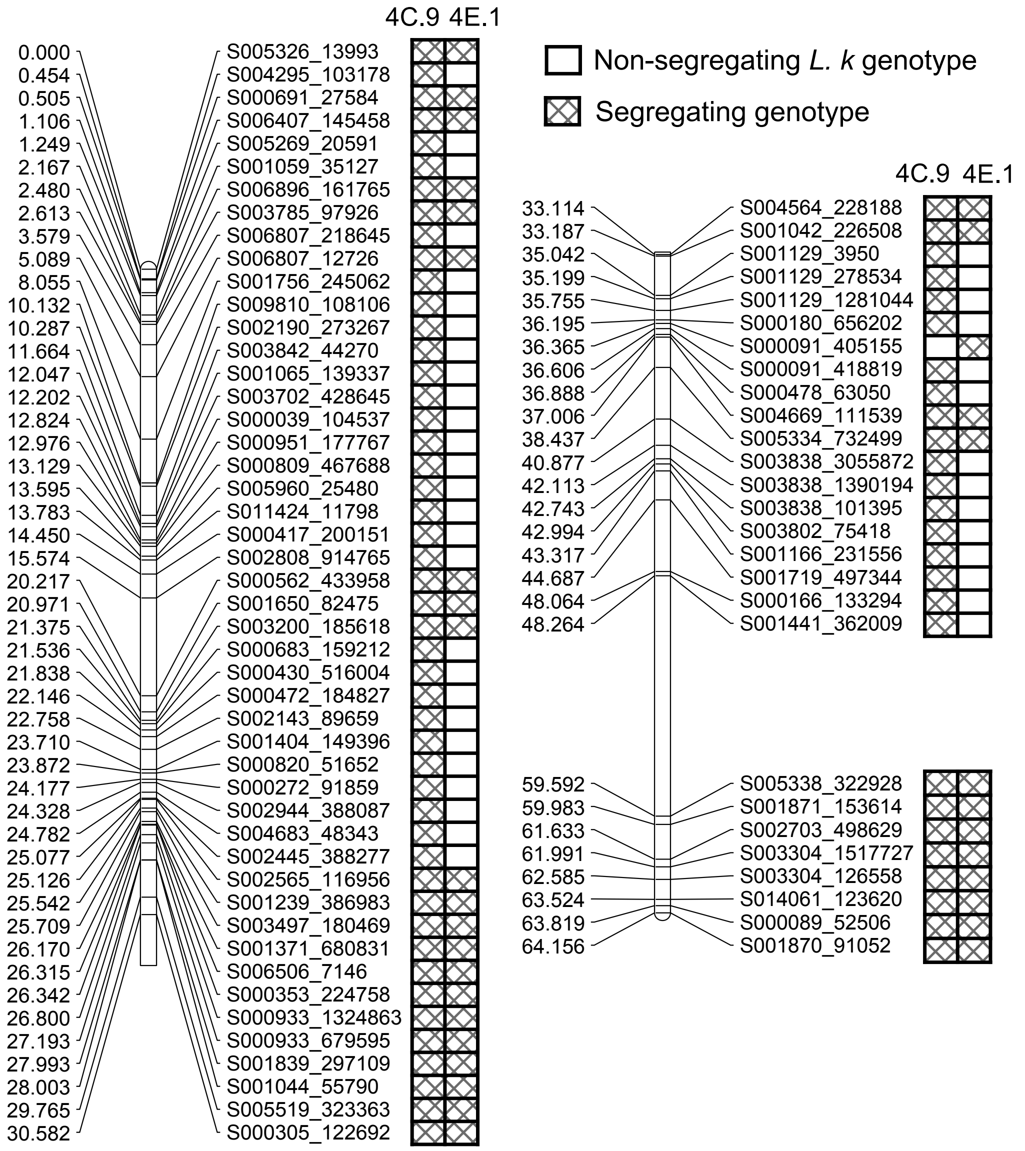


Figure S1. Segregation pattern of markers on the integrated linkage map in the two $F_2$ mapping families 4C.9 and 4E.1.

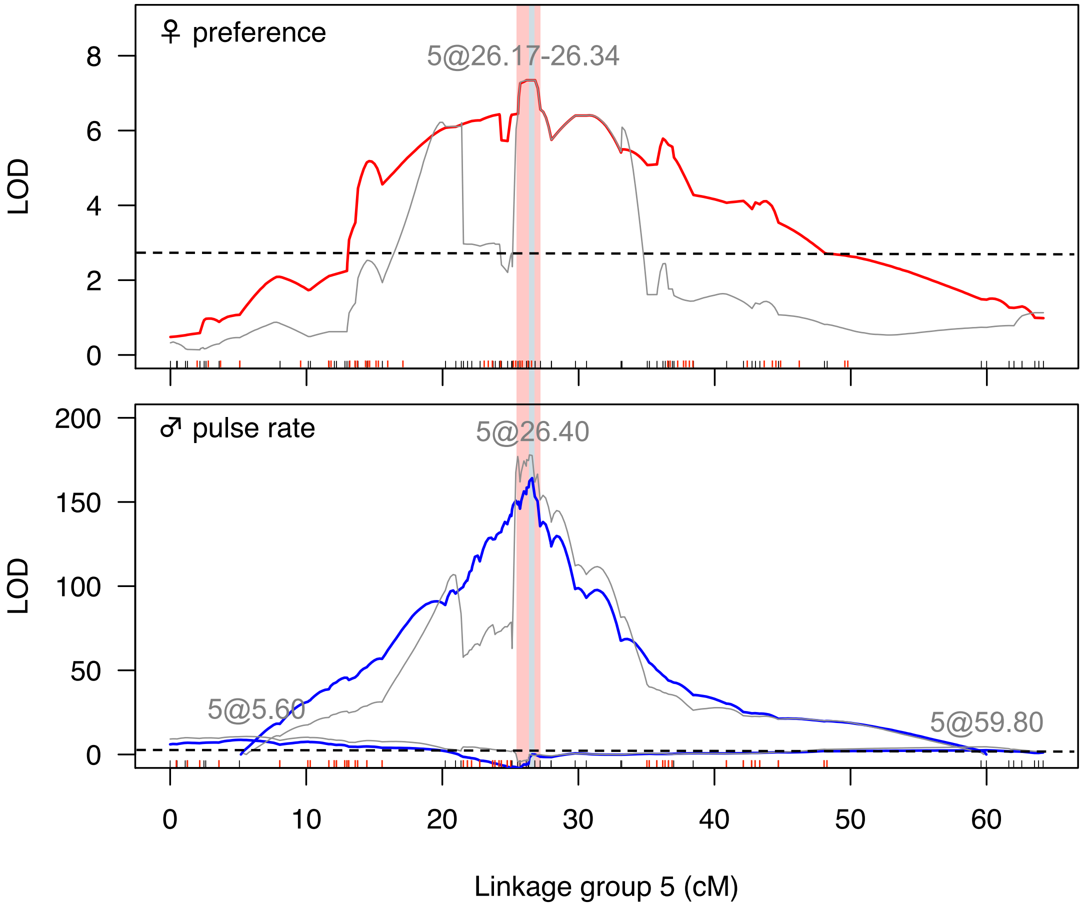


Figure S2. LOD profiles from the final multiple QTL mapping (MQM) models for interspecific variation in male pulse rate and female peak preference for pulse rate. LOD profiles from MQM models using only genotypes of marker loci segregating in a 1:2:1 ratio (i.e., genotypes at marker loci segregating in only one family were treated as missing data, and thus simulated, in the non-segregating family) are shown in red (female preference) and blue (male pulse rate). LOD profiles from MQM models using all known genotypes, including those not segregating in one of the two families, are shown in grey. Markers in red indicate those segregating in only one of the two families. The red and blue shaded areas indicate 1.5-LOD support confidence intervals for preference and pulse rate respectively.
